# Super-resolution vibrational imaging based on photoswitchable Raman probe

**DOI:** 10.1101/2022.08.28.505494

**Authors:** Jingwen Shou, Ayumi Komazawa, Yuusaku Wachi, Minoru Kawatani, Hiroyoshi Fujioka, Spencer John Spratt, Takaha Mizuguchi, Kenichi Oguchi, Fumiaki Obata, Ryo Tachibana, Yoshihiro Misawa, Ryosuke Kojima, Yasuteru Urano, Mako Kamiya, Yasuyuki Ozeki

**Author notes:** These authors contributed equally: Jingwen Shou, Ayumi Komazawa.

## Abstract

Super-resolution vibrational microscopy is a promising tool to increase the degree of multiplexing of nanometer-scale biological imaging, because the spectral linewidth of molecular vibration is about 50 times narrower than that of fluorescence. However, current techniques of super-resolution vibrational microscopy still suffer from various limitations including the need for cell fixation, high power loading or complicated frequency-modulated detection schemes. Herein we utilize photoswitchable stimulated Raman scattering (SRS) to develop a method that we call reversible saturable optical Raman transitions (RESORT) microscopy, which overcomes these limitations. We first describe a new kind of photoswitchable Raman probe designated DAE620 and then we employ a standard SRS detection scheme to validate its signal activation and depletion characteristics when exposed to low-power (microwatt level) continuous-wave laser light. By harnessing the SRS signal depletion of DAE620 through a donut-shaped beam, we demonstrate super-resolution vibrational imaging of mammalian cells with excellent chemical specificity and spatial resolution beyond the optical diffraction limit. Our results indicate RESORT microscopy to be an effective tool with high potential for multiplexed super-resolution imaging of live cells.

---

Far-field super-resolution fluorescence microscopy has enabled the visualization of intracellular structures in biological systems with nanometer-scale resolution, overcoming the diffraction limit of light microscopy^1–2^. Furthermore, techniques of multicolor super-resolution fluorescence imaging have been developed^3–4^, though it is difficult to greatly increase the number of resolvable colors due to the large bandwidth of the fluorescence signals^5^. In contrast, Raman scattering, which probes molecular vibrational states, can generate sharp spectral signatures with about 50 times narrower linewidth than that of fluorescence^6–8^. Hence, extending Raman microscopy beyond the diffraction limit seems a promising approach to increase the degree of multiplexing for far-field super-resolution light microscopy.

Among recent advances in coherent anti-Stokes Raman scattering (CARS) microscopy and stimulated Raman scattering (SRS) microscopy^9–12^, several works of high-resolution or super-resolution vibrational imaging have been reported, though these still have various limitations^13–24^. The method of generating a smaller diffraction-limited point spread function (PSF) by shortening the wavelength of SRS light from the near infrared region to the visible region^1^ is associated with significant cytotoxicity. Also, the combination of expansion microscopy with SRS detection^14–16^ is not compatible with biological imaging of live samples. As regards methods based on high-order nonlinearity of Raman scattering processes^17–23^, extremely high instantaneous light power is required, which can cause severe photodamage. Also, when a shorter pulse time duration is exploited to provide higher instantaneous power, the broader spectral bandwidth derived from the Fourier transform limit decreases the spectral resolution of vibrational imaging. In regard to the method employing the combination of stimulated Raman excited fluorescence (SREF) and stimulated emission depletion (STED)^24^, a frequency-modulated scheme of SREF excitation and detection is needed to obtain sufficient signal depletion, and this increases the complexity of the system. In addition, efficient SREF excitation requires critical matching between the CARS wavelength of the two-color pulse laser system and the absorption wavelength of the probe molecule, which increases the difficulty of molecular design of the probes.

Herein, we propose reversible saturable optical Raman transitions (RESORT) microscopy as a new kind of optical far-field super-resolution Raman microscopy which can realize vibrational imaging beyond the optical diffraction limit with low-intensity photoswitching lasers and a relatively simple system configuration. We chose SRS as the basic Raman imaging technology, as it can provide high vibrational contrast of intracellular structures without a non-resonant background^9–12^. Then, we developed a novel diarylethene-based Raman probe named DAE620, the SRS signal of which in the fingerprint region can be activated or depleted sufficiently and directly by additional low-power continuous-wave (CW) laser light. The photoswitchable Raman response of DAE620 shows high fatigue resistance and a fast photoswitching speed. Based on the effective SRS signal depletion of DAE620 by a donut-shaped beam, we show that RESORT microscopy of HeLa cells provides vibrational biological contrasts with excellent chemical specificity and high spatial resolution beyond the optical diffraction limit. We believe that RESORT microscopy promises to be a useful supplementary tool for achieving highly multiplexed super-resolution imaging of biological systems, especially living cells.

## RESULTS

### RESORT microscopy based on photoswitchable SRS

Among the methods of far-field super-resolution fluorescence microscopy^1–2^, the direct regulation of PSF is most compatible with SRS imaging, considering the sensitivity and nonlinear excitation of SRS. Distinct from STED microscopy, reversible saturable optical fluorescence transitions (RESOLFT) microscopy based on photoswitchable fluorescence probes enables PSF modification under low power^25^. Recently, photoswitchable SRS spectroscopy and microscopy^26–28^ have been demonstrated by exploiting photochromic molecules that can be photoswitched between two distinct isomers by applying ultraviolet (UV) or visible light, turning the Raman signal on or off. Inspired by RESOLFT microscopy, we propose the concept of RESORT microscopy, in which an additional CW laser light including a donut-shaped beam is introduced to regulate the SRS detection of a photoswitchable Raman probe (Fig. 1).

**Figure 1.**
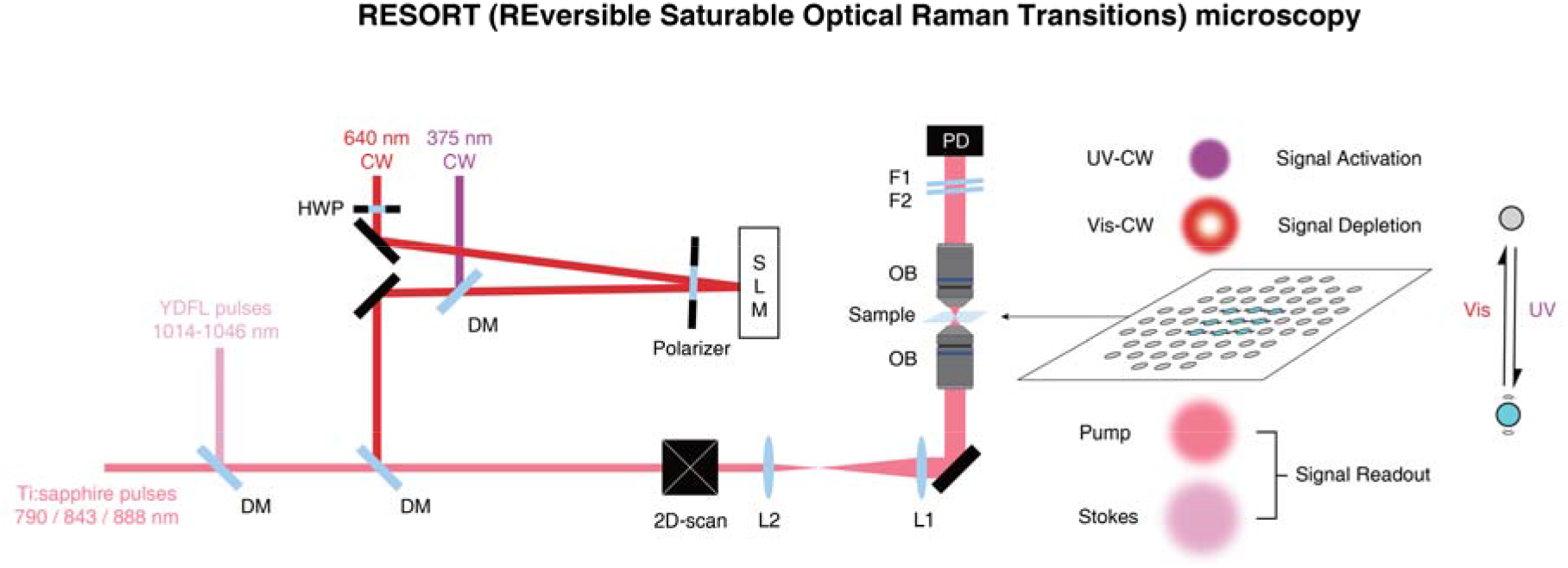
RESORT microscopy for super-resolution vibrational imaging. CW, continuous-wave laser; HWP, half wave plate; SLM, spatial light modulator; DM, dichroic mirror; YDFL, ytterbium-doped fiber laser system; L, lens; OB, objective; F, filter; PD, photodiode; UV, ultraviolet; Vis, visible.

Specifically, picosecond narrowband pump and Stokes pulses of SRS are provided by a Ti:sapphire laser (Coherent, Mira 900D) and a home-built ytterbium-doped fiber laser system, respectively, for the SRS signal readout. A UV laser and a visible laser provide CW light at 375 nm and 640 nm, respectively, for the signal on/off control of the photoswitchable Raman probe. In particular, the 640 nm visible CW light is reflected by a spatial light modulator (SLM) at a small incidence angle to introduce phase modulation. A half wave plate (HWP) and a polarizer are used to make the polarization of 640 nm light linear and the angle of polarization matched with the axis of SLM, guaranteeing the sufficient phase modulation depth by SLM. Then, all laser beams are combined by dichroic mirrors and led to a point-by-point scanning microscope composed of broadband achromatic optical elements and water-immersion objective lenses (Olympus, N.A. = 1.2). After passing through the sample, the pump pulses are filtered out from the transmitted light and detected to generate SRS signals.

For RESORT imaging, azimuthal 2π phase modulation by SLM is applied to generate a donut-shaped focal spot of 640-nm CW light at the sample plane. The smaller beam size of UV CW light compared to the size of the entrance pupil of the focusing objective is utilized to make the focal spot size of UV light comparable with the size of the PSF of normal SRS. After the competition of signal activation by UV light and signal depletion by visible light, only the photoswitchable molecules at the very center of the donut-shaped beam focus can maintain the on-state of the Raman signal and be detected by SRS. As a result, PSF of RESORT imaging can possess a narrower full width at half maximum (FWHM) than that of normal SRS imaging, enabling super-resolution vibrational imaging beyond the optical diffraction limit.

### Photophysical and SRS properties of existing photoswitchable dyes

Next, we set out to develop a photoswitchable Raman probe suitable for RESOLFT microscopy based on the diarylethene (DAE) scaffold^29^, a representative photochromic dye, as it has recently been reported that some DAE derivatives show changes in their SRS spectra upon photoirradiation^26,27^. For this purpose, we started by examining the effect of absorption wavelength on the SRS intensity of the closed form by comparing commercially available DAE derivatives, 1,2-dicyano-1,2-bis(2,4,5-trimethyl-3-thienyl)ethene (CMTE) and 1,2-bis(2,4-dimethyl-5-phenyl-3-thienyl)-3,3,4,4,5,5-hexafluoro-1-cyclopentene (DMPT-PFCP), whose closed forms have maximum absorption wavelengths at 531 nm and 579 nm, respectively (Supplementary Fig. 1a,b). These DAEs exhibit photoswitching between the open form with UV absorption and the closed form with visible absorption, accompanied with distinct changes in the SRS spectra (Supplementary Fig. 1c). According to the results of molecular orbital calculation, the SRS peaks of CMTE at 1496 cm^-1^ and DMPT-PFCP at 1493 cm^-1^ were assigned to stretching across the thiophene and indole rings (Supplementary Fig. 2, Supplementary Movies 1–2). The SRS signal intensity ratio between the open/closed forms at the above-mentioned wavenumbers was only 4- to 5-fold for either CMTE or DMPT-PFCP. Interestingly, the signal intensity of DMPT-PFCP with absorption at longer wavelength was higher than that of CMTE; this can be presumably explained by the electronic pre-resonance (epr) effect^6^ (Supplementary Fig. 1d). As a higher signal-to-noise ratio (SNR) is crucial for RESORT microscopy to achieve a high spatial resolution and fast image acquisition, we next set out to develop a photoswitchable Raman probe with a higher SRS signal intensity by extending the absorption wavelength.

### Design of new photoswitchable Raman probe for RESORT imaging

Considering the near-infrared (NIR) pump light used in our SRS system^30,31^, we speculated that it would be possible to obtain a sufficiently strong SRS signal enhancement due to the epr effect when the molecular absorption of the closed form resides in the red region. Among the previously reported DAE derivatives, we focused on asymmetric DAEs consisting of a thiophene ring at one side and an indole ring at the other side as a candidate scaffold with molecular absorption in the red region and superior fatigue resistance performance^29^ (Supplementary Fig. 3). By modifying a reported asymmetric DAE so that the closed form does not show absorption over 840 nm which would lead to undesirable one-photon absorption by the SRS light, we newly synthesized an asymmetric DAE designated as DAE620 (Fig. 2a). The open form of DAE620 exhibits absorption in the UV region with a maximum wavelength at 354 nm. Upon UV light irradiation, the open form switches to the closed form with the appearance of red absorption with a maximum wavelength at 623 nm (Fig. 2b).

**Figure 2.**
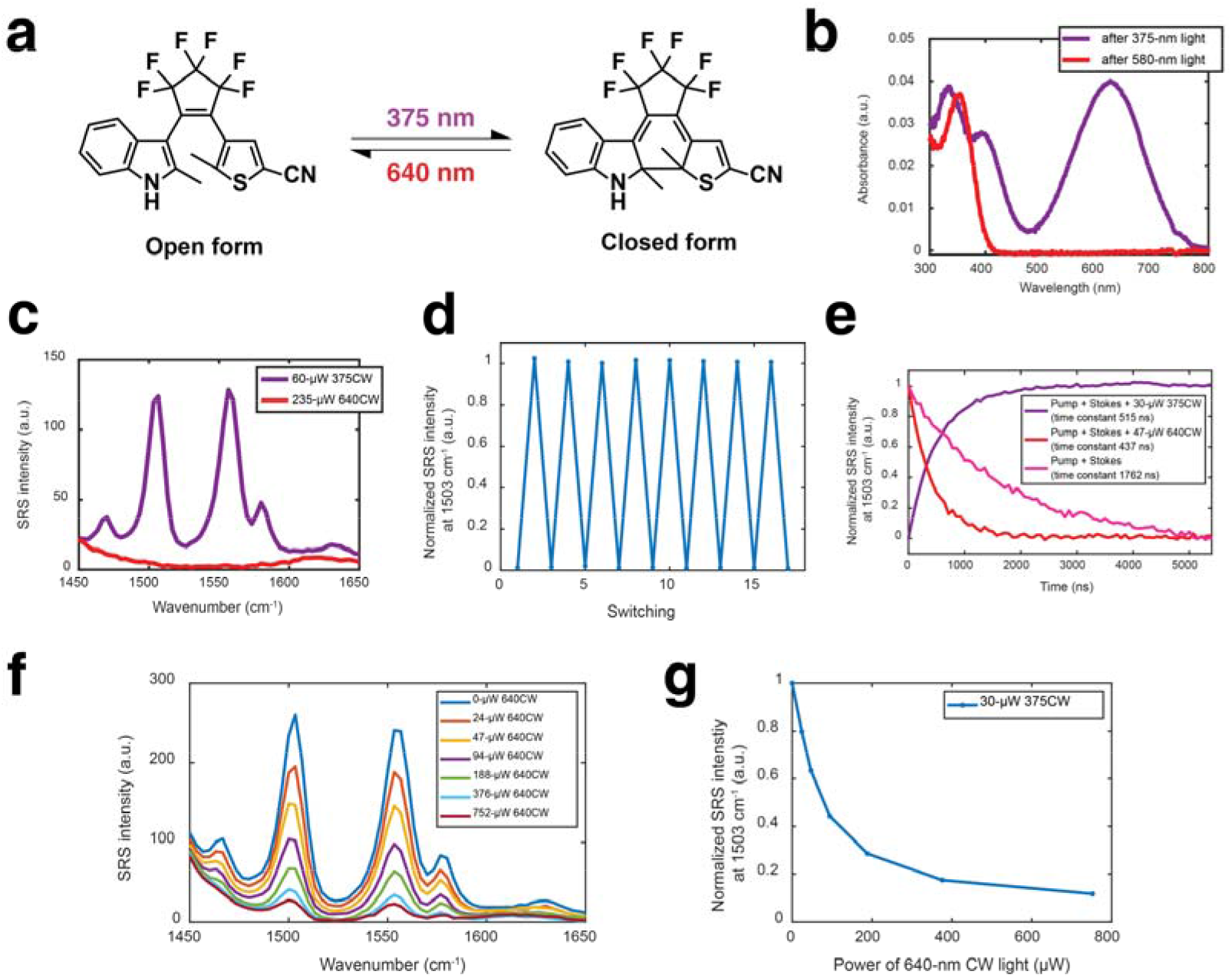
In vitro photoswitching properties of DAE620. **a** Structures of the open and closed forms of DAE620. **b** Absorption spectra of 10 μM DAE620 in DMSO under different conditions of CW light irradiation. **c** SRS spectra of 10 mM DAE620 in DMSO under different conditions of CW light irradiation. Note that the residual spectral background at the low wavenumber side derives from the solvent, DMSO. **d** Evaluation of photoswitching fatigue of the SRS signal of 10 mM DAE620 in DMSO. **e** Evaluation of photoswitching speed of 10-mM DAE620 in DMSO. **f** SRS spectra of 10 mM DAE620 in DMSO under irradiation with 30 μW 375 nm CW light and different powers of 640 nm CW light. **g** Normalized SRS intensity at 1503 cm^-1^ extracted from **f**.

Next, we measured the SRS spectrum of DAE620 (Fig. 2c). Under 375 nm CW light irradiation, the closed form of DAE620 exhibits two strong peaks at 1503 cm^-1^ and 1557 cm^-1^, two medium peaks at 1470 cm^-1^ and 1580 cm^-1^, and a small peak at 1630 cm^-1^ in the fingerprint region. Based on molecular orbital calculations, the two representative peaks at 1503 cm^-1^ and 1557 cm^-1^ were assigned to stretching of the thiophene ring and stretching across the thiophene and indole rings, respectively (Supplementary Fig. 2, Supplementary Movies 3–4). The linewidth of the sharp Raman peak at 1503 cm^-1^ is evaluated as about 15 cm^-1^, showing high chemical specificity, which is beneficial for multiplexing. Under 640 nm CW light irradiation, the DAE620 molecules all switch to open form and the five Raman peaks mentioned above disappear. The SRS signal intensity at 1503 cm^-1^ exhibits more than 25-fold change upon photoswitching, demonstrating that sufficient control of the Raman signal is achieved by the additional CW light. The relative Raman intensity versus EdU (RIE) value^32^ of the closed form of DAE620 was calculated to be 6.35, which is higher than those of CMTE (RIE: 1.43) and DMPT-PFCP (RIE: 2.95) (Supplementary Fig. 1d, Supplementary Table 1). The higher SRS signal intensity of DAE620 is considered to be a consequence of the epr effect due to the longer absorption wavelength. We also examined the SRS spectrum of DAE620 in the silent region, as it has nitrile group in its structure (Supplementary Fig. 4). As expected, the closed form of DAE620 exhibits a single sharp peak at 2217 cm^-1^, but with lower signal intensity than that at 1503 cm^-1^ in the fingerprint region. When comparing the SRS signal intensity between the open and closed forms, the signal activation upon photoswitching was limited to 3-fold probably owing to the limited electronic-vibrational coupling strength of the C≡N triple bond. Besides, either the closed or open form of DAE exhibits a vibration-independent spectral background in the silent region, which could be problematic for potential multiplexing. Thus, we focused on the peak at 1503 cm^-1^ of DAE620 as the target peak for the following experiments, considering the signal intensity of two isomers and the spectral background at the Raman off-resonance frequency.

### Photoswitching kinetics of DAE620 under SRS microscopy

In order to verify the compatibility of DAE620 with RESORT imaging, we firstly confirmed the photoswitching fatigue resistance of DAE620 (Fig. 2d, Supplementary Fig. 5). Both the SRS signal intensity and the absorbance of two isomers were preserved over multiple cycles. Further, we measured the photoswitching speed of DAE620 (Fig. 2e). DAE620 molecules were firstly placed in a photostationary state by applying 375 nm or 640 nm CW light, and then were photoswitched by immediately switching the condition of light irradiation. The time constant (*τ*) of photoswitching by 30 μW 375 nm CW light was evaluated as *τ* = 515 ns, and that by 47 μW 640 nm CW light was evaluated as *τ* = 437 ns. Such fast photoswitching (*τ* < 1 μs) under low-power CW light irradiation is highly compatible with a point-scan microscope. Although we observed that two-photon absorption by pump and Stokes pulses also caused photoconversion from the closed form to the open form with disappearance of the SRS signal, as previously reported^27^, the time constant of photoswitching in the absence of CW light was evaluated as *τ* = 1762 ns (Fig. 2e), indicating that the photoswitching by one-photon absorption is dominant even when the CW light power is very low.

Considering that the competition between signal activation by UV light and signal depletion by visible light is directly related to the signal output in RESORT imaging, we measure the SRS spectra of DAE620 when 375 nm and 640 nm CW lights were simultaneously applied (Fig. 3c). When the 375 nm CW light power was set constant as 30 μW, the peak intensity at 1503 cm^-1^ was gradually depleted by applying higher-power 640 nm CW light (Fig. 2f,g). Meanwhile, no additional spectral background appeared in the fingerprint region. The normalized SRS signal intensity reached 0.44 when 30 μW 375 nm and 94 μW 640 nm CW lights were applied. Eventually when 752 μW 640 nm light was applied, the normalized SRS signal intensity reached an almost saturated value of 0.11, a little higher than that with only 640 nm CW light. These results demonstrate that sufficient Raman signal depletion can be directly realized through photoswitching under microwatt-level CW power, without the need for high instantaneous light power or a complicated frequency-modulated detection scheme.

**Figure 3.**
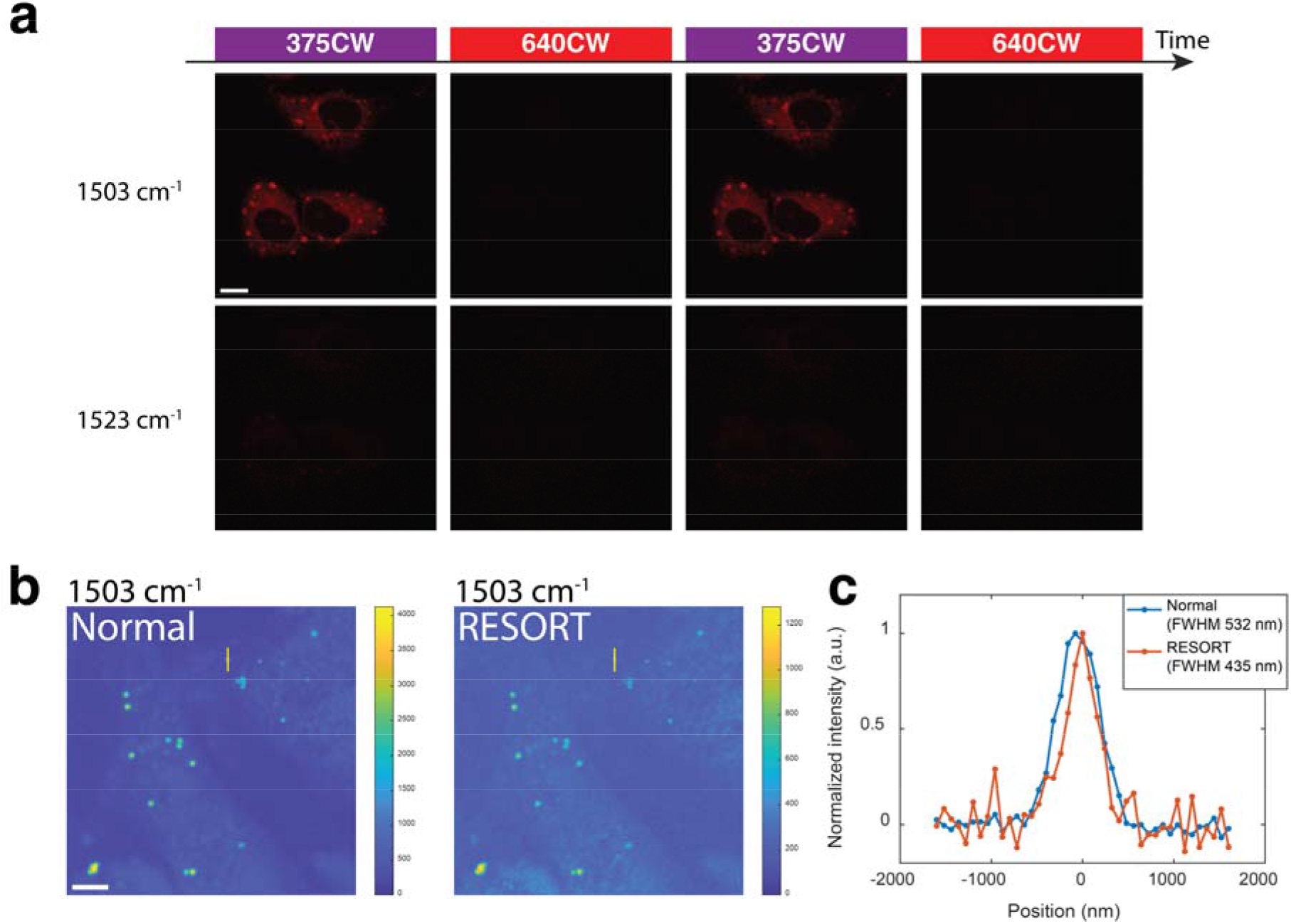
Cellular imaging of HeLa cells stained with DAE620. **a** Photoswitchable SRS imaging of fixed HeLa cells stained with DAE620. Scale bar, 10 μm (**a**), 5 μm (**b**). **b** Normal SRS imaging and RESORT imaging of fixed HeLa cells. **c** Cross section along yellow lines in **b**. Pump: 39 mW, Stokes: 52 mW, 375CW: 14 μW, 640CW: 92 μW, pixel dwell time: 227 μs (**b**-**c**).

### RESORT imaging with DAE620 in cells

To confirm whether the photoswitching properties of DAE620 are preserved in cells, we performed SRS imaging of HeLa cells stained with DAE620 (Fig. 3a). The spectral characterization of the stained cells shows results similar to those of the *in vitro* measurement (Supplementary Fig. 6), despite some additional spectral background derived from cross-phase modulation (XPM) and intrinsic Raman signals of cells in the fingerprint region. When applying 375 nm light, we observed significant SRS signal activation at the Raman on-resonance wavenumber (1503 cm^-1^), but not at the Raman off-resonance wavenumber (1523 cm^-1^). In contrast, when 640 nm CW light was applied, the SRS signal was switched off. These photoswitching cycles were repeatable. These results suggested that the photoswitching properties and chemical specificity of DAE620 are well preserved even in the intracellular environment. Further, we confirmed *ex vivo* applicability of DAE620 by applying it to *Drosophila* fat body tissue. We observed that DAE620 tends to localize to lipid droplets in the fat body tissue. Again, the SRS signal of DAE620 was successfully photoswitched by alternately applying 375 nm and 640 nm CW light (Supplementary Fig. 7). These results indicate the potential utility of DAE620 for RESORT imaging of live tissues.

Next, to demonstrate the effectiveness of RESORT microscopy, we observed fixed HeLa cells stained with DAE620, which showed small aggregates inside the cells. In the case of normal SRS imaging where only 375 nm CW light is applied in addition to pump and Stokes light, the FWHM of a small aggregate was 532 nm (Fig. 3b). Considering the diffraction-limited resolution of 342 nm when using 888 nm pump light, we can calculate the object FWHM as approximately 408 nm. In contrast, when an additional donut-shaped beam of 640 nm CW light was applied for signal depletion, the FWHM of the same region by RESORT imaging was 435 nm. By means of deconvolution calculation, we can evaluate the spatial resolution of RESORT microscopy as approximately 151 nm (see Methods), indicating two-fold improvement of spatial resolution beyond the optical diffraction limit. In another field of view (FOV), we find three aggregates distributed very closely (Supplementary Fig. 8). By exploiting RESORT imaging, each of them is observed with sharper edges and they can be distinguished from each other more clearly thanks to the improved PSF.

### RESORT imaging of mitochondria in cells by DAE620-Mito

In order to showcase the ability of RESORT to observe important subcellular organelles^33–34^, we focused on mitochondria as a target structure and introduced triphenylphosphonium (Mito-tag) into DAE620 via a butyl linker to prepare DAE620-Mito (Fig. 4a). Despite the slightly longer maximum absorption wavelength of the closed form of DAE620-Mito (644 nm) than that of DAE620 (623 nm) due to *N*-alkylation of DAE620, the SRS measurement of DAE620-Mito shows good consistency with the results obtained using DAE620 both *in vitro* and *in cellulo* (Supplementary Fig. 9, Supplementary Fig. 10). Further, we confirmed that the SRS signal of DAE620-Mito colocalized well with the fluorescence signal of MitoTracker Green, verifying the mitochondrial targeting of DAE620-Mito (Fig. 4b).

**Figure 4.**
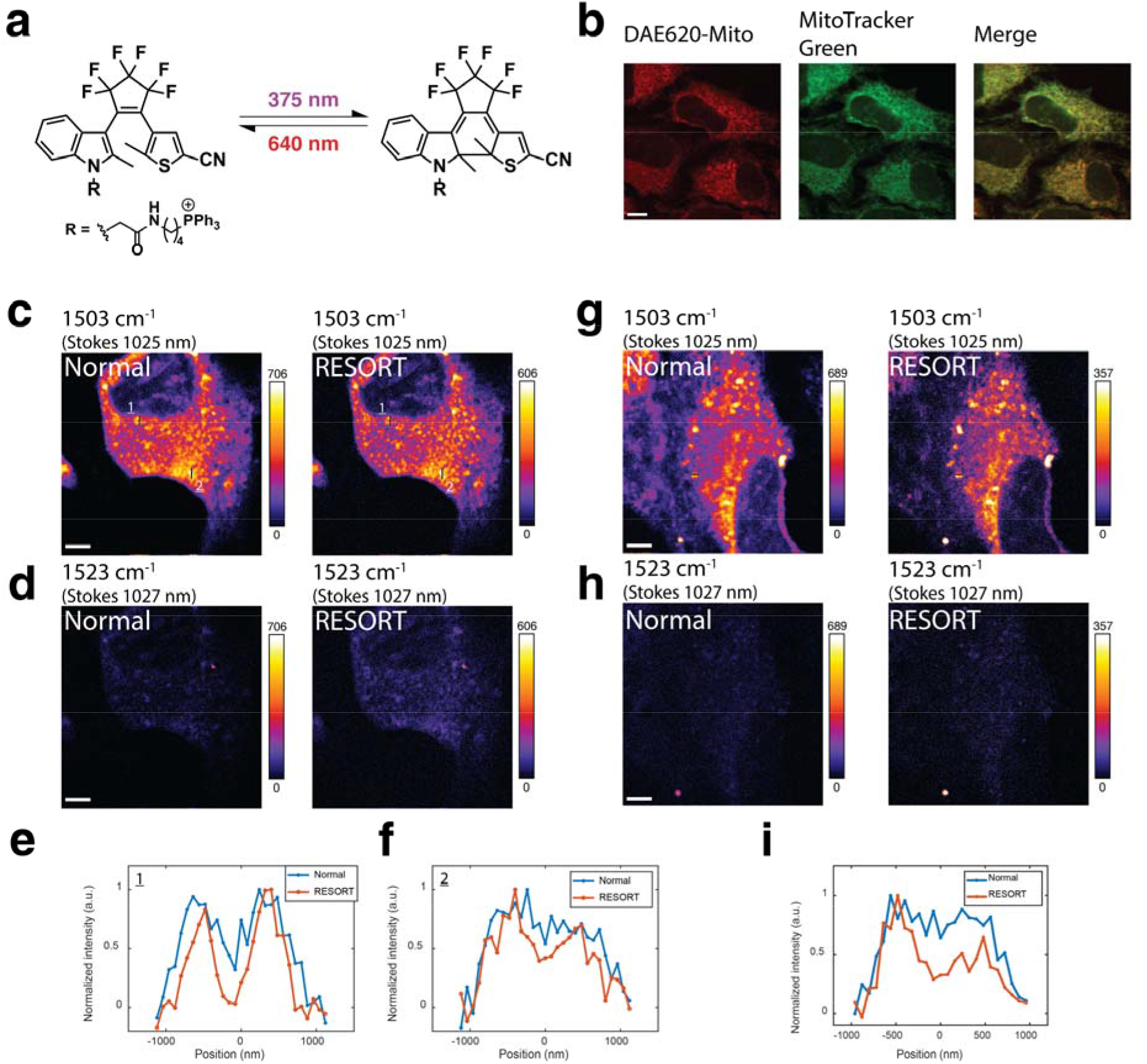
Cellular imaging of HeLa cells stained with DAE620-Mito. **a** Structures of the open and closed forms of DAE620-Mito. **b** Colocalization imaging of DAE620-Mito (SRS imaging at 1503 cm^-1^) and MitoTracker Green (fluorescence imaging with 488 nm excitation and 510 nm detection) for live HeLa cells. **c-d** Normal SRS imaging and RESORT imaging of fixed HeLa cells at 1503 cm^-1^ (Raman on-resonance) (**c**) and 1523 cm^-1^ (Raman off-resonance) (**d**). **e-f** Cross sections along black lines in position 1 (**e**) and position 2 (**f**) in **c**. **g-h** Normal SRS imaging and RESORT imaging of live HeLa cells at 1503 cm^-1^ (Raman on-resonance) (**g**) and 1523 cm^-1^ (Raman off-resonance) (**h**). **i** Cross sections along black lines in **g**. Scale bars, 10 μm (**b**), 5 μm (**c-d**, **g-h**). Pump: 39 mW, Stokes: 52 mW, 375CW: 14 μW, 640CW: 184 μW, pixel dwell time: 227 μs (**c**-**f**). Pump: 26 mW, Stokes: 52 mW, 375CW: 14 μW, 640CW: 230 μW, pixel dwell time: 227 μs (**g**-**i**).

Then, we performed RESORT imaging of fixed HeLa cells stained with DAE620-Mito (Fig. 4c-f). With normal SRS imaging, the two ends of a folded mitochondrion appear fused (Fig. 4c,e, position 1). In contrast, with the improved spatial resolution of RESORT imaging, the two ends are seen to be separated, providing a more accurate characterization of the organelle organization. In another position in the same FOV, normal SRS imaging under the diffraction limit gives a flat-topped mitochondrial profile without detailed textures (Fig. 4c,f, position 2). In contrast, RESORT imaging breaks the diffraction limit and shows detailed contrast with a lower signal intensity. These results show that super-resolution vibrational imaging by RESORT can reveal rich nanoscale information beyond the diffraction limit in biological samples. As a control, when imaging at a Raman off-resonance wavenumber by tuning the Stokes wavelength only 2 nm (corresponding to 20 cm^-1^ wavenumber tuning), no clear contrast was observed due to chemical specificity of vibrational imaging (Fig. 4d). The results suggest that super-resolution vibrational imaging by RESORT microscopy preserves the advantage of the narrow linewidth of Raman signatures, which will be helpful for achieving a high degree of multiplexing.

Encouraged by these results, we further tried live-cell RESORT imaging (Fig. 4g-i). In normal SRS imaging, two mitochondria look connected (Fig. 4g,i). In contrast, RESORT imaging clearly revealed a gap between these mitochondria, suggesting that the significant improvement of spatial resolution by RESORT is maintained in live-cell imaging. Similar to the case in imaging of fixed cells, no clear contrast can be distinguished at a Raman off-resonance wavenumber (Fig. 4h), again supporting the chemical specificity. As RESORT imaging can be done by applying low-power CW light, it should be beneficial by reducing potential damage due to phototoxicity during super-resolution vibrational imaging of live cells.

## DISCUSSION

Inspired by the RESOLFT method, we propose RESORT microscopy as a new type of super-resolution Raman microscopy, which modifies the PSF of SRS based on photoswitchable Raman probes. We synthesized a novel photoswitchable Raman probe DAE620, whose Raman signal can be controlled sufficiently and directly by introducing an additional microwatt-level CW laser light. Further, by targeting DAE620 to mitochondria and harnessing the photoswitching-based signal activation/depletion, we succeeded in the super-resolution vibrational imaging of mitochondria in HeLa cells by RESORT microscopy with high chemical specificity and a spatial resolution two-fold beyond the diffraction limit.

Compared to the combination of expansion microscopy with SRS detection^14–16^, RESORT does not require cell fixation and processing, and thus is compatible with live-cell imaging. Compared to some super-resolution methods based on high-order nonlinearity of Raman scattering processes^17–23^, which enable label-free super-resolution imaging, RESORT imaging relies on the labelling of photoswitchable probes, which permit the observation of specific subcellular targets. Advantageously, signal depletion based on photoswitchable probes in RESORT can be achieved with a low-power CW light at microwatt level, which reduces the potential phototoxicity and thus should be more compatible with live cells. As for the method combining SREF and STED^24^, which has greater detection sensitivity than our RESORT, the signal depletion of SREF requires a complicated frequency-modulated detection scheme. In contrast, RESORT can achieve sufficient signal depletion directly with the general SRS system configuration. In addition, the photoswitching can occur as long as the CW laser lines match the absorption spectrum of the photoswitchable molecules, and does not require critical wavelength matching, which is necessary for SREF excitation.

Since the linewidth of Raman scattering is generally about 50 times narrower than that of fluorescence excitation and emission, RESORT microscopy offers high chemical specificity with the potential for highly multiplexed super-resolution imaging. Compared to the reported photoswitchable Raman probes based on DAEs, DAE620 shows a much lower residual signal of the open form, which would again be beneficial for potential multiplexed RESORT imaging considering that not all molecules are converted to the closed form upon UV light irradiation (Supplementary Fig. 12). In order to achieve multiplexed super-resolution vibrational imaging, we need to expand the availability of photoswitchable Raman probes that show SRS signal activation/depletion at different wavenumbers. Although the photoswitching response of the nitrile triple bond of DAE620 in the silent region is limited, we should be able to expand the pallet in the silent region if we can optimize the electronic-vibrational coupling strength of the nitrile bond by appropriate chemical modification. Alternatively, it remains possible to achieve multiplexing in the fingerprint region^6^ by exploiting and optimizing other kinds of photoswitchable molecules with different scaffold structures. With such strategies to expand the range of photoswitchable vibrational palettes, we anticipate that RESORT microscopy will open up a wide variety of applications, including super-multiplex biological imaging.

## METHODS

### SRS microscope

A Ti:sapphire laser (Coherent, Mira900D) provides picosecond narrowband pump pulses at 76 MHz repetition rate. The central wavelength of the pump pulses is fixed at 790 nm for imaging in the C-H stretching region (2800–3100 cm^-1^), at 843.26 nm for imaging in the silent region (2000–2300 cm^-1^), or at 888.2 nm for imaging in the fingerprint region (1400–1700 cm^-1^). The Stokes pulses for SRS imaging are provided by a laser system working at a repetition rate of 38 MHz. Specifically, a home-built polarization-maintaining ytterbium-doped fiber laser (PM-YDFL) followed by spectral broadening fiber optics generates broadband seed pulses. These broadband pulses are introduced into a tunable optical bandpass filter. After tunable spectral filtering, the narrowband pulses are further amplified by a two-stage polarization-maintaining ytterbium-doped fiber amplifier (PM-YDFA). As a result, the central wavelength of the YDFA output can be tuned from 1014 nm to 1046 nm, corresponding to a wavenumber range of 300 cm^-1^. Fractions of pump light and Stokes light are extracted and launched to a two-photon absorption photodiode, the signal of which is input to a feedback circuit to control the cavity of YDFL, so that the pump laser and the Stokes laser can be sub-harmonically synchronized^31^. The pump light and Stokes light are spatially combined by a dichroic mirror and temporally aligned by adjusting the time delay line, and finally led to an x-y scanner. Two types of x-y scanners are used by switching the optical path with flipping mirrors. One, consisting of a resonant galvanometric scanner and an ordinary galvanometric scanner, is used to provide two-dimensional scanning with a fixed pixel dwell time of 71 ns^30,31^. The other, consisting of two ordinary galvanometric scanners, is used to provide two-dimensional scanning with an adjustable and relatively longer pixel dwell time. The plane of the scanners is imaged onto the pupil of a water-immersion objective lens (Olympus, 60×, N.A.=1.2). The light is focused on the specimen and the transmitted light is collected by another objective lens (Olympus, 60×, N.A.=1.2). The pump light is filtered out from the transmitted light and detected by a Si photodiode, the output of which is further demodulated by a lock-in amplifier at 38 MHz to generate the SRS signal.

We evaluated the spatial resolution of normal SRS imaging by imaging 300-nm diameter polystyrene beads at a wavenumber of 3050 cm^-1^ (Supplementary Fig. 11). The FWHM of the contrast of a single bead is 378 nm. Through the calculation of deconvolution based on the Gaussian assumption described below, we can evaluate the spatial resolution of normal SRS imaging as approximately 347 nm, close to theoretical diffraction-limited resolution of 319 nm.

### Fluorescence microscope

A 488 nm CW laser is used as the light source for fluorescence excitation. A point-by-point scan microscope including a x-y scanner and a water-immersion objective lens (Olympus, 60×, N.A.=1.2) is used to image the sample. The epi-fluorescence is spatially filtered by a confocal pinhole and spectrally filtered to remove laser light. Then the fluorescence is introduced into a tunable optical bandpass filter and finally detected by a photomultiplier to generate the fluorescence signal^8^.

### Calculation of spatial resolution

The theoretical spatial resolution of normal SRS imaging can be calculated by applying the following equation,

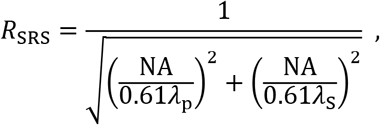

where NA is the numerical aperture of the objective; *λ_p_* is the wavelength of pump light; *λ_s_* is the wavelength of Stokes light; R_SRS_ is the theoretical spatial resolution of normal SRS imaging. Based on the assumption of a Gaussian profile, the FWHM of the real object R_obj_ can be calculated according to the following equation,

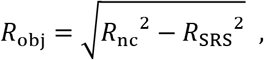

where *R_nc_* is the FWHM of the contrast imaged by normal SRS microscopy; R_SRS_ is the spatial resolution of normal SRS imaging. Based on the assumption of a Gaussian profile, the spatial resolution of RESORT imaging can be calculated by means of the following equation,

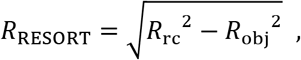

where *R_rc_* is the FWHM of the contrast imaged by RESORT microscopy; R_RESORT_ is the spatial resolution of RESORT imaging.

### Cell preparation for imaging

HeLa cells provided from RIKEN BRC were used for analysis. The cells were cultured on glass coverslips in a 4-well plate containing Dulbecco’s modified Eagle’s medium (DMEM) (Invitrogen, 12320032) supplemented with 10% fetal bovine serum (FBS) (GE Health Care Biosciences, SH30079.03). All cell cultures were maintained in a humidified environment at 37°C and 5% CO_2_.

#### Photoswitchable SRS imaging of fixed HeLa cells stained with DAE620

The cells were stained with 60 μM DAE620 for 3 hours in the culture medium. Then the cells were fixed in fixation buffer (BioLegend, 420801) for 10 minutes. Finally, the glass coverslip was washed with PBS and assembled into a chamber using an imaging spacer filled with PBS for imaging.

#### Photoswitchable SRS imaging of fixed HeLa cells stained with DAE620-Mito

The cells were stained with 58 μM DAE620-Mito for 4 hours in the culture medium. Then the cells were fixed in fixation buffer for 10 minutes. Finally, the glass coverslip was washed with PBS and assembled into a chamber using an imaging spacer filled with PBS for imaging.

#### RESORT imaging of fixed HeLa cells stained with DAE620

The cells were stained with 60 μM DAE620 for 3 hours in the culture medium. Then the cells were fixed in fixation buffer for 10 minutes. Finally, the glass coverslip was washed with PBS and assembled into a chamber using an imaging spacer filled with PBS for imaging.

#### RESORT imaging of fixed HeLa cells stained with DAE620-Mito

The cells were stained with 29 μM DAE620-Mito for 3 hours in the culture media. Then the cells were fixed in fixation buffer for 10 minutes. Finally, the glass coverslip was washed with PBS and assembled into a chamber using an imaging spacer filled with PBS for imaging.

#### RESORT imaging of live HeLa cells stained with DAE620-Mito

The cells were stained with 29 μM DAE620-Mito for 4 hours in the culture medium. Then, the glass coverslip was washed with PBS and assembled into a chamber using an imaging spacer filled with PBS for imaging.

#### Colocalization imaging of live HeLa cells stained with DAE620-Mito and MitoTracker Green

The cells were stained with 29 μM DAE620-Mito for 4 hours and 100 nM MitoTracker Green (Invitrogen, M7514) for 30 minutes in the culture medium. Then, the glass coverslip was washed with PBS and assembled into a chamber using an imaging spacer filled with PBS for imaging.

### Tissue preparation

*Drosophila melanogaster* Canton-S fruit flies were maintained on a standard yeast-based diet containing 4% yeast (Asahi), 6% glucose (Nisshoku), 4.5% cornflour (Nippon) and 0.8% agar with preservatives (0.4% propionic acid (Wako, 163-04726) and 0.03% butyl p-hydroxybenzoate (Wako, 028-03685)). Fat bodies were dissected from third-instar larvae, mounted in a 35 mm glass-bottomed dish, and cultured in S2 medium containing 300 μM DAE620 for 3.5 hours. Then, the fat bodies were assembled into a chamber using an imaging spacer filled with PBS for imaging.

## Acknowledgments

This work was supported by JSPS KAKENHI Grant Number JP20H05724, JP20H05725, JP20H05726, JP19K22242, JP20H02650, JP22H02193, and JP19J22546, by JSPS Core-to-Core Program, A. Advanced Research Networks, by JST CREST JPMJCR1872, by The Naito Foundation (to M.K.), by The Mitsubishi Foundation (to M.K.), by Daiichi Sankyo Foundation of Life Science (to M.K.), and by Quantum Leap Flagship Program of MEXT JPMXS0118067246. J.S. is supported by International Research Fellow of the Japan Society for the Promotion of Science. We thank G. Dai for his technical support on fiber laser system.

## Author contributions

J.S. and Y.W. built the imaging system, and K.O. developed the data acquisition codes. A.K. H.F. and Y.M. synthesized the photochromic compounds, J.S., A.K., M. Kawatani, S.S. and T.M. analyzed the photochemical properties of the compounds. F.O. and S.S. prepared *Drosophila* samples. R.T. performed quantum chemical calculations. J.S., H.F. A.K., M. Kawatani and Y.W. performed imaging experiments. J.S. and A.K. conducted data analysis. R.K. and Y.U. supervised the research. M. Kamiya. and Y.O. planned and supervised the entire research. All authors participated in writing the manuscript.

## Data availability

The data that support the findings of this study are available from the corresponding authors upon reasonable request.

## Additional information

Supplementary information is available for this paper. Reprints and permission information is available at www.nature.com/reprints.

## References

1. Sahl, S., Hell, S. & Jakobs, S. Fluorescence nanoscopy in cell biology. Nat. Rev. Mol. Cell Biol. 18, 685–701 (2017).

2. Sigal, Y., Zhou, R. & Zhuang, X. Visualizing and discovering cellular structures with super-resolution microscopy. Science. 361, 880–887 (2018).

3. Bates, M., Huang, B., Dempsey, G. & Zhuang, X. Multicolor super-resolution imaging with photo-switchable fluorescent probes. Science. 317, 1749–1753 (2007).

4. Shechtman, Y., Weiss, L., Backer, A., Lee, M. & Moerner, W. Multicolour localization microscopy by point-spread-function engineering. Nat. Photonics. 10, 590–594 (2016).

5. Lakowicz, J. Principles of Fluorescence Spectroscopy 3rd edn. (Springer, New York, 2006).

6. Wei, L., Chen, Z., Shi, L., Long, R., Anzalone, A., Zhang, L., Hu, F., Yuste, R., Cornish, V. & Min, W. Super-multiplex vibrational imaging. Nature. 544, 465–470 (2017).

7. Hu, F., Zeng, C., Long, R., Miao, Y., Wei, L., Xu, Q. & Min, W. Supermultiplexed optical imaging and barcoding with engineered polyynes. Nat. Methods. 15, 194–200 (2018).

8. Shou, J., Oda, R., Hu, F., Karasawa, K., Nuriya, M., Yasui, M., Shiramizu, B., Min, W. & Ozeki, Y. Super-multiplex imaging of cellular dynamics and heterogeneity by integrated stimulated Raman and fluorescence microscopy. iScience. 24, 102832 (2021).

9. Hu, F., Shi, L. & Min, W. Biological imaging of chemical bonds by stimulated Raman scattering microscopy. Nat. Methods. 16, 830–842 (2019).

10. Min, W., Freudiger, C., Lu, S. & Xie, X. Coherent nonlinear optical imaging: Beyond fluorescence microscopy. Annu. Rev. Phys. Chem. 62, 507–530 (2011).

11. Cheng, J.-X. & Xie, X. Vibrational spectroscopic imaging of living systems: An emerging platform for biology and medicine. Science. 350, aaa8870 (2015).

12. Camp Jr, C. & Cicerone, M. Chemically sensitive bioimaging with coherent Raman scattering. Nat. Photonics. 9, 295–305 (2015).

13. Bi, Y., Yang, C., Chen, Y., Yan, S., Yang, G., Wu, Y., Zhang, G. & Wang, P. Near-resonance enhanced label-free stimulated Raman scattering microscopy with spatial resolution near 130 nm. Light Sci. Appl. 7, 81 (2018).

14. Qian, C., Miao, K., Lin, L., Chen, X., Du, J. & Wei, L. Super-resolution label-free volumetric vibrational imaging. Nat. Commun.,. 12, 3648 (2021).

15. Shi, L., Klimas, A., Gallagher, B., Cheng, Z., Fu, F., Wijesekara, P., Miao, Y., Ren, X., Zhao, Y. & Min, W. Super-resolution vibrational imaging using expansion stimulated Raman scattering microscopy. Adv. Sci. 9, 2200315 (2022).

16. Miao, K., Lin, L., Qian, C. & Wei, L. Label-free super-resolution imaging enabled by vibrational imaging of swelled tissue and analysis. J. Vis. Exp. 183, e63824 (2022). doi:10.3791/63824

17. Yonemaru, Y., Palonpon, A., Kawano, S., Smith, N., Kawata, S. & Fujita, K. Super-spatial-and-spectral-resolution in vibrational imaging via saturated coherent anti-Stokes Raman scattering. Phys. Rev. Appl. 4, 014010 (2015).

18. Gong, L., Zheng, W., Ma, Y. & Huang, Z. Saturated stimulated-Raman-scattering microscopy for far-field superresolution vibrational imaging. Phys. Rev. Appl.,. 11, 034041 (2019).

19. Cleff, C., Groß, P., Fallnich, C., Offerhaus, H., Herek, J., Kruse, K., Beeker, W., Lee, C. & Boller, K. Ground-state depletion for subdiffraction-limited spatial resolution in coherent anti-Stokes Raman scattering microscopy. Phys. Rev. A. 86, 023825 (2012).

20. Silva, W., Graefe, C. & Frontiera, R. Toward label-free super-resolution microscopy. ACS Photon. 3, 79–86 (2015).

21. Kim, D., Choi, D., Kwon, J., Shim, S., Rhee, H. & Cho, M. Selective suppression of stimulated Raman scattering with another competing stimulated Raman scattering. J. Phys. Chem. Lett. 8, 6118–6123 (2017).

22. Würthwein, T., Irwin, N. & Fallnich, C. Saturated Raman scattering for sub-diffraction-limited imaging. J. Chem. Phys. 151, 194201 (2019).

23. Gong, L., Zheng, W., Ma, Y. & Huang, Z. Higher-order coherent anti-Stokes Raman scattering microscopy realizes label-free super-resolution vibrational imaging. Nat. Photonics. 14, 115–122 (2019).

24. Xiong, H., Qian, N., Miao, Y., Zhao, Z., Chen, C. & Min, W., Super-resolution vibrational microscopy by stimulated Raman excited fluorescence. Light Sci. Appl. 10, 87 (2021).

25. Kwon, J., Hwang, J., Park, J., Han, G., Han, K. & Kim, S. RESOLFT nanoscopy with photoswitchable organic fluorophores. Sci. Rep. 5, 17804 (2015).

26. Shou, J. & Ozeki, Y., 2021. Photoswitchable stimulated Raman scattering spectroscopy and microscopy. Opt. Lett. 46, 2176–2179 (2021).

27. Ao, J., Fang, X., Miao, X., Ling, J., Kang, H., Park, S., Wu, C. & Ji, M. Switchable stimulated Raman scattering microscopy with photochromic vibrational probes. Nat. Commun. 12, 3089 (2021).

28. Lee, D., Qian, C., Wang, H., Li, L., Miao, K., Du, J., Shcherbakova, D., Verkhusha, V., Wang, L. & Wei, L. Toward photoswitchable electronic pre-resonance stimulated Raman probes. J. Chem. Phys. 154, 135102 (2021).

29. Irie, M. Diarylethenes for memories and switches. Chem. Rev. 100, 1685–1716 (2000).

30. Ozeki, Y., Umemura, W., Otsuka, Y., Satoh, S., Hashimoto, H., Sumimura, K., Nishizawa, N., Fukui, K. & Itoh, K. High-speed molecular spectral imaging of tissue with stimulated Raman scattering. Nat. Photonics. 6, 845–851 (2012).

31. Ozeki, Y., Asai, T., Shou, J. & Yoshimi, H. Multicolor stimulated Raman scattering microscopy with fast wavelength-tunable Yb fiber laser. IEEE J. Sel. Top. Quantum Electron. 25, 7100211 (2019).

32. Yamakoshi, H., Dodo, K., Palonpon, A., Ando, J., Fujita, K., Kawata, S. & Sodeoka, M. Alkyne-tag Raman imaging for visualization of mobile small molecules in live cells. J. Am. Chem. Soc. 134, 20681–20689 (2012).

33. Wong, Y., Ysselstein, D. & Krainc, D. Mitochondria–lysosome contacts regulate mitochondrial fission via RAB7 GTP hydrolysis. Nature. 554, 382–386 (2018).

34. Han, Y., Li, M., Qiu, F., Zhang, M. & Zhang, Y. Cell-permeable organic fluorescent probes for live-cell long-term super-resolution imaging reveal lysosome-mitochondrion interactions. Nat. Commun. 8, 1307 (2017).

