## Supplementary Information for "Super-resolution vibrational imaging based on photoswitchable Raman probe"

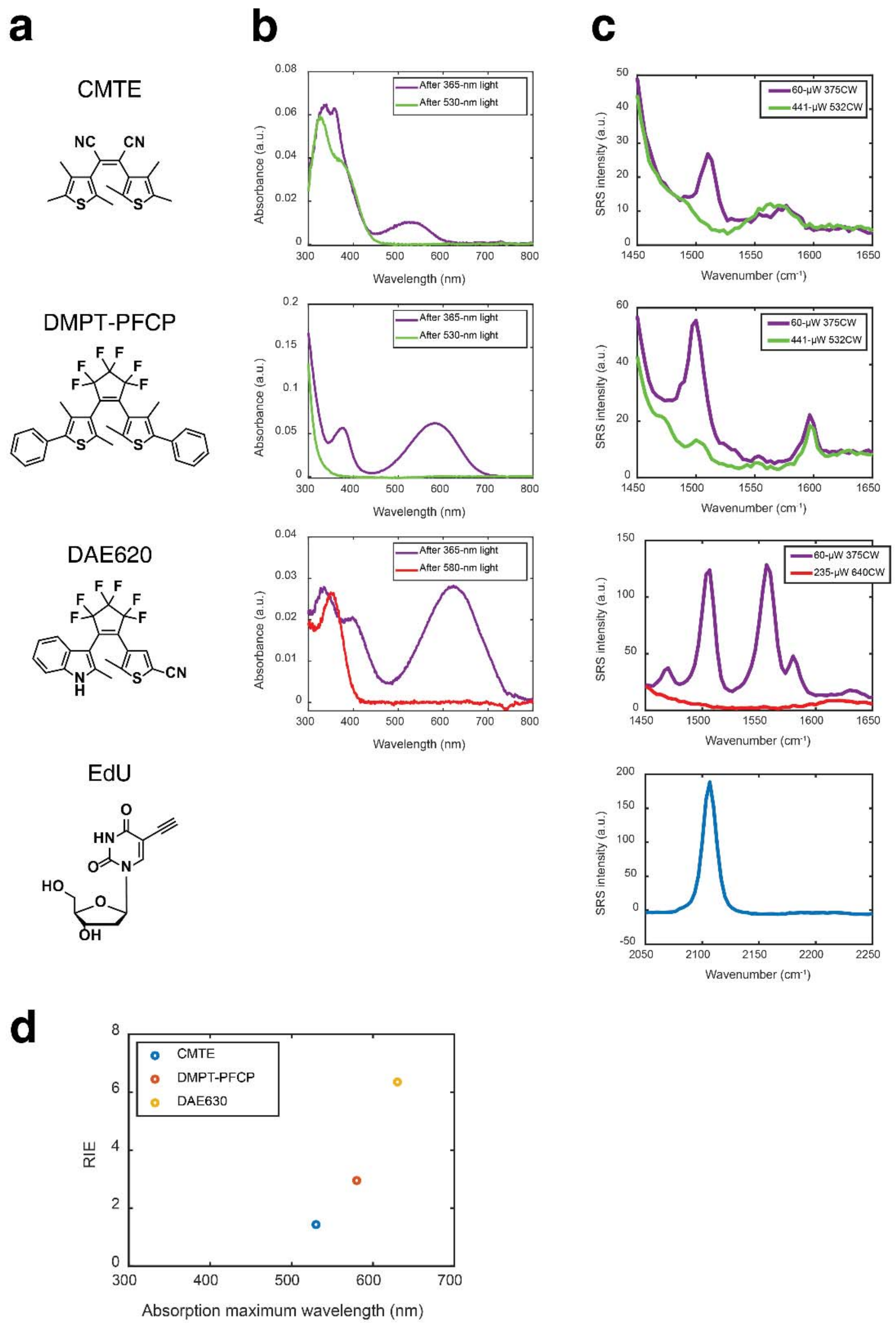


**Supplementary Figure 1 | a** Molecular structures of CMTE, DMPT-PFCP, DAE620 and EdU. **b** Absorption spectra of 10 μM CMTE, 10 μM DMPT-PFCP and 7 μM DAE620 in DMSO. **c** SRS spectra of 10 mM CMTE, 10 mM DMPT-PFCP, 10 mM DAE620 and 100 mM EdU in DMSO. **d** Comparison of RIE of different photoswitchable molecules. CMTE, RIE 1.43, 1510 $cm^{-1}$; DMPT-PFCP, RIE 2.95, 1500 $cm^{-1}$; DAE620, RIE 6.35, 1503 $cm^{-1}$.

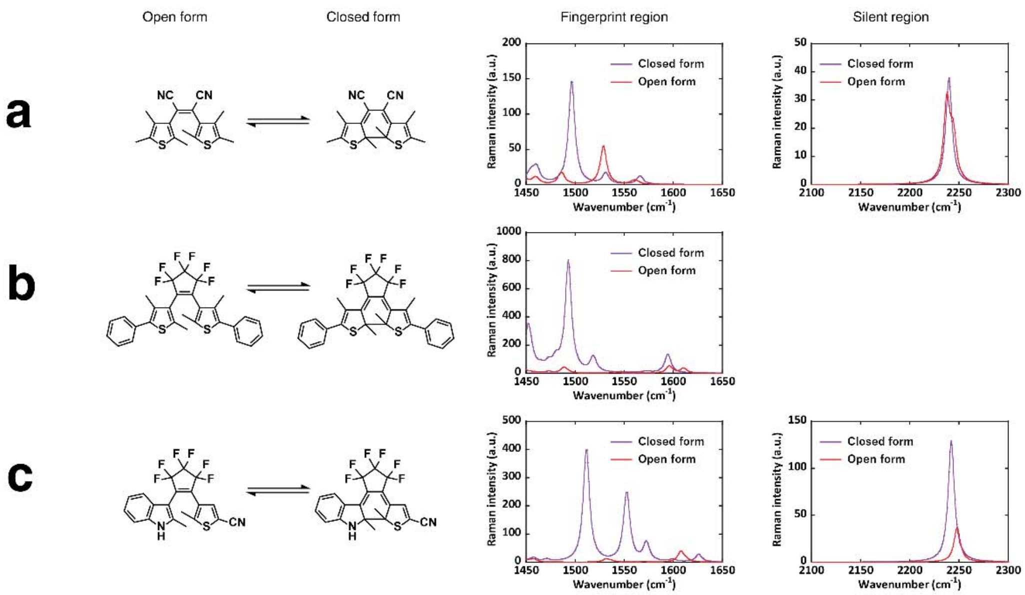


**Supplementary Figure 2 |** Molecular structures and simulation of Raman responses of CMTE (**a**), DMPT-PFCP (**b**), and DAE620 (**c**).

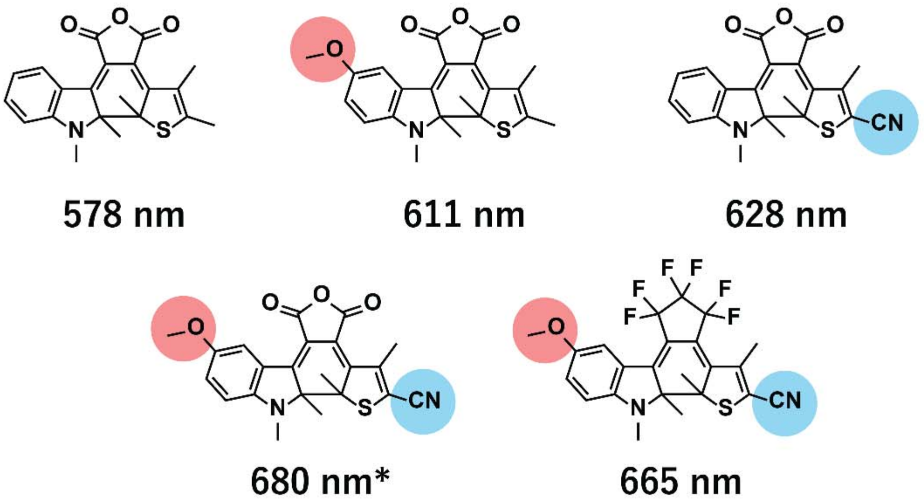


**Supplementary Figure 3 |** Previously reported asymmetric DAEs composed of a thiophene ring and an indole ring[1]. *According to the literature[1], the absorption edge of the closed-ring isomer extends to 860 nm, meaning that one-photon absorption by our near-infrared pump light can occur with derivatives having both electron-donating and -withdrawing substituents.

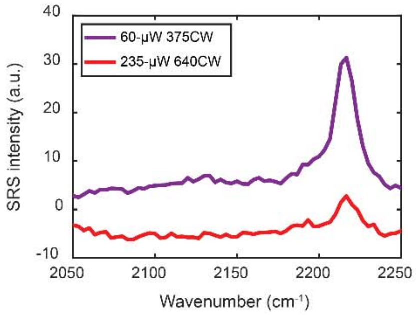


**Supplementary Figure 4 |** SRS spectra of 10 mM DAE620 in DMSO in the silent region under different conditions of CW light irradiation.

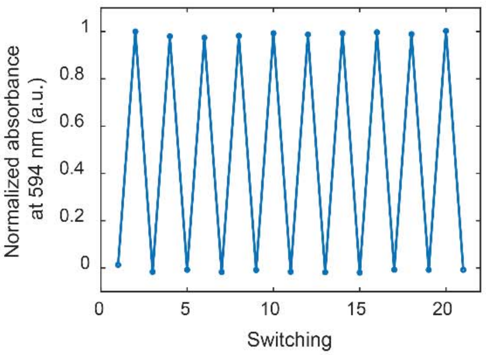


**Supplementary Figure 5 |** Evaluation of photoswitching fatigue of absorbance of 10 μM DAE620 in acetonitrile under repetitive 365 nm and 580 nm CW light irradiation.

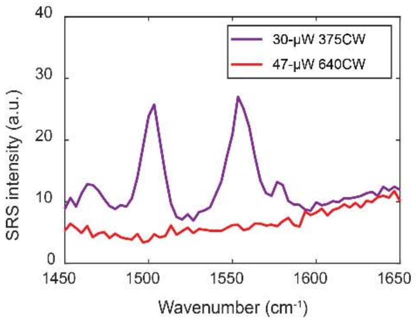


**Supplementary Figure 6 |** Photoswitchable SRS spectra of fixed HeLa cells stained with DAE620.

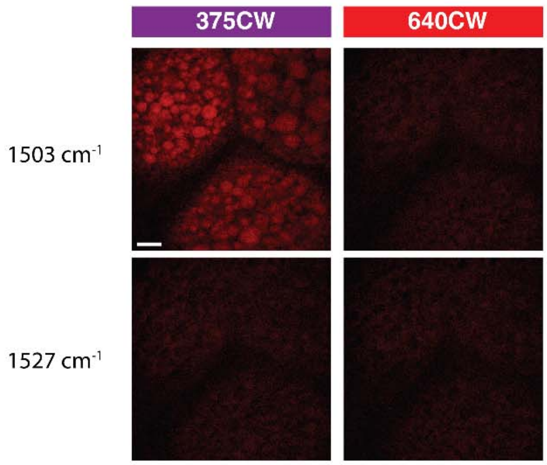


**Supplementary Figure 7 |** Photoswitchable SRS imaging of *Drosophila* larval fat bodies stained with DAE620. Scale bar, 10 µm.

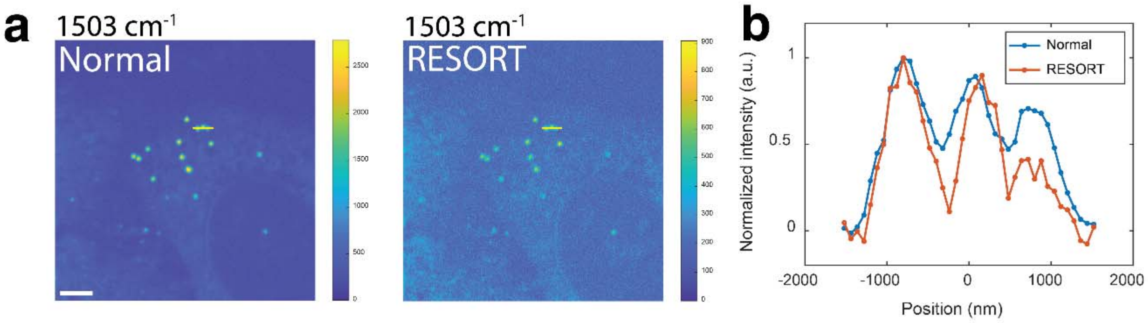


**Supplementary Figure 8 | a** Normal SRS imaging and RESORT imaging of fixed HeLa cells. **b** Cross sections along yellow lines in **a**. Pump: 39 mW, Stokes: 52 mW, 375CW: 14 µW, 640CW: 92 µW, pixel dwell time: 227 µs.

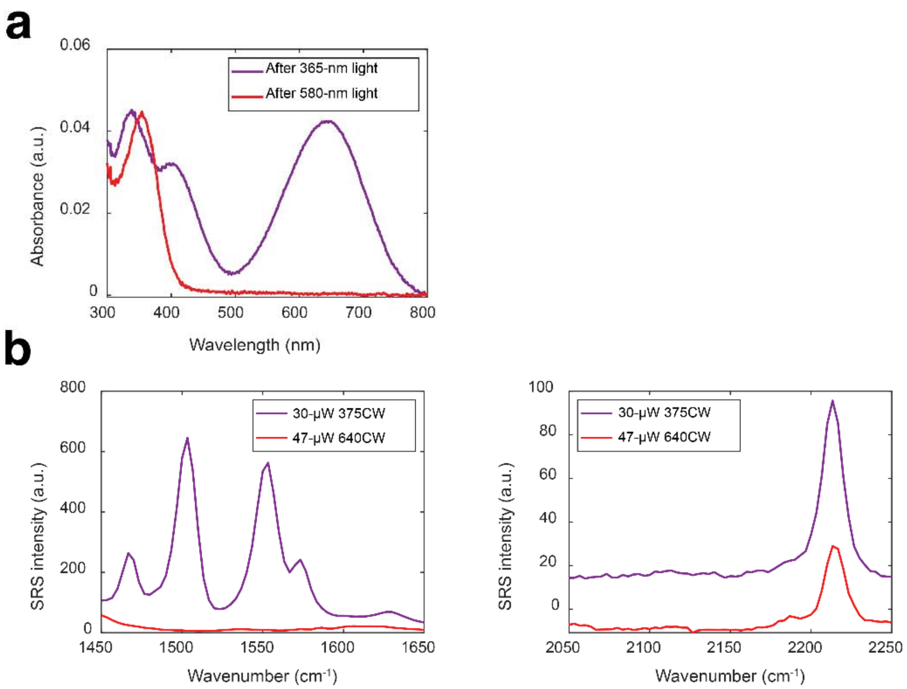


**Supplementary Figure 9 | a** Absorption spectra of 10 μM DAE620-Mito in DMSO under different conditions of CW light irradiation. **b** SRS spectra of 29 mM DAE620-Mito in DMSO under different conditions of CW light irradiation.

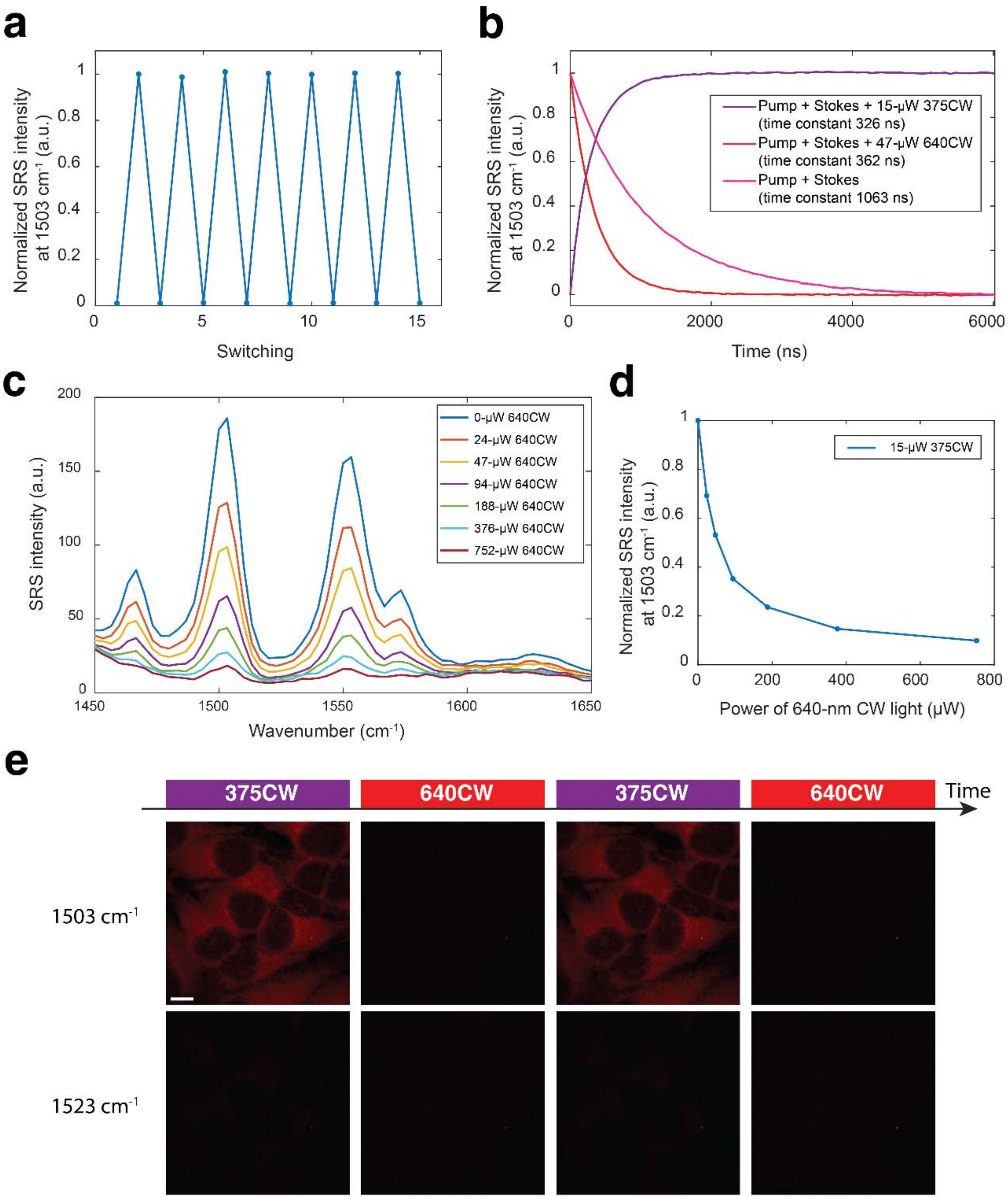


**Supplementary Figure 10 | Characterizations of DAE620-Mito. a** Evaluation of photoswitching fatigue of SRS intensity of 29 mM DAE620-Mito in DMSO. **b** Evaluation of photoswitching speed of 29 mM DAE620-Mito in DMSO. **c** SRS spectra of 29 mM DAE620-Mito in DMSO under irradiation with 15 µW 375 nm CW light and different power levels of 640 nm CW light. **d** Normalized SRS intensity at 1503 $cm^{-1}$ extracted from **c**. **e** Photoswitchable SRS imaging of fixed HeLa cells stained by DAE620-Mito. Scale bar, 10 µm (**e**).

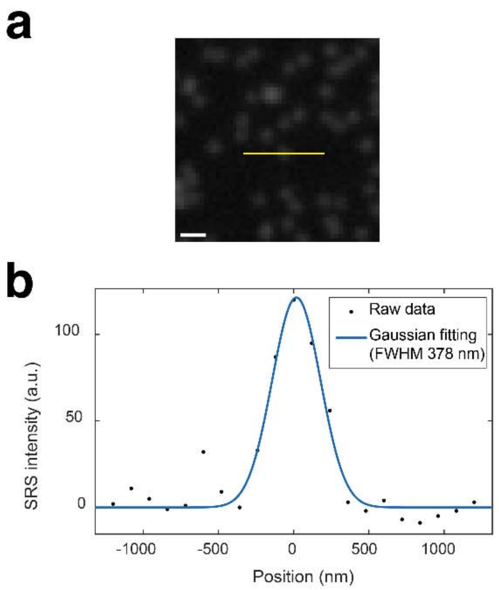


**Supplementary Figure 11 | a** Normal SRS imaging of polystyrene microsphere (300 nm diameter) suspension in distilled water at 3050 $cm^{-1}$. **b** Cross section along yellow line in **a** and corresponding Gaussian fitting. Scale bar, 0.75 μm (**a**).

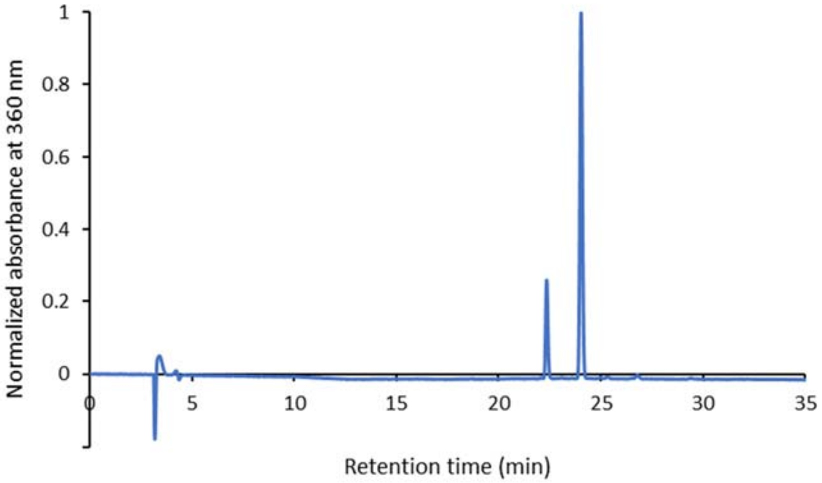


**Supplementary Figure 12 |** Determination of PSS composition (open/closed) of DAE620 by analytical HPLC. A 15 µM solution of DAE620 in DMSO was irradiated with 365 nm light until the absorbance at 627 nm reached a plateau. Then, the PSS composition was determined by analytical HPLC. Absorbance at 360 nm, which is the isosbestic point of the absorption spectra of the open and closed forms, was monitored. The open/closed components were calculated to amount to 20% and 80%, respectively. PSS was calculated from the area percentage of each peak.

| | DAE620 | DMPT-PFCP | CMTE |
|---|---|---|---|
| Absorption max (nm) | 623 | 579 | 531 |
| RIE | 6.35 | 2.95 | 1.43 |
| Peak wavenumber ($cm^{-1}$) | 1503 | 1500 | 1510 |

**Supplementary Table 1 |** Comparison of absorption maximum, RIE, and peak wavenumber of different photoswitchable molecules after UV light irradiation.

### Supplementary Note

#### Materials.

General chemicals were of the best grade available, supplied by Tokyo Chemical Industries, Wako Pure Chemical or Aldrich Chemical Company, and were used without further purification. Dimethyl sulfoxide (DMSO, fluorometric grade) for the spectrometric measurements was purchased from Dojindo.

#### Instruments.

Purification by column chromatography was performed on a YFLC-AI580 chromatograph (Yamazen, Osaka, Japan). Preparative HPLC was performed on an HPLC system composed of reverse-phase columns of Inertsil ODS-3 10 mm × 250 mm or 20 mm × 250 mm (GL Sciences, Tokyo, Japan), with a pump (JASCO, PU-2080 or PU-2087) and a detector (JASCO, MD-2018). $^{1}H$ NMR and $^{13}C$ NMR spectra were recorded on a Bruker AVANCEIII400 instrument (400 MHz for $^{1}H$, 101 MHz for $^{13}C$) with chemical shifts (δ) given in ppm relative to residual solvents for $^{1}H$ and $^{13}C$. High-resolution mass spectra were recorded on a Bruker micrOTOF II, using electrospray ionization (ESI). All experiments were carried out at 298 K, unless otherwise specified.

#### Raman Spectrum Calculation

The structure optimization and spectrum calculation were performed with Gaussian09[2], at the level of B3LYP/6-31G (d) without any consideration of symmetricity. The wavenumber of the Raman spectrum is scaled by 0.9614, according to the literature[3].

#### Absorption measurements.

Absorption spectra were obtained with a UV-2450 UV/Vis spectrometer (Shimadzu). Probes were dissolved in DMSO (fluorometric grade, Dojindo) to obtain stock solutions. For DAE620 photoirradiation experiments *in vitro*, DAE620 solutions in DMSO in a quartz cuvette were illuminated at 580 nm (10.1 mW/cm$^2$) using a Xe lamp (Asahi Spectra Inc., MAX-303) equipped with a 581.00 ± 5.00 nm bandpass filter, with stirring at room temperature, while CMTE and DMPT-PFCP solutions were illuminated at 530 nm (10.6 mW/cm$^2$) using the same lamp equipped with a 530.50 ± 5.50 nm bandpass filter, with stirring at room temperature. Ultraviolet (UV) irradiation was performed at 365 nm (0.66 mW/cm$^2$) with a handy UV lamp (AS ONE, SLUV-4) for all compounds.

#### Purity analysis by HPLC.

In order to confirm the purity of DAE620 and DAE620-Mito, each dye solution was analyzed with an HPLC system composed of reverse-phase columns of Inertsil ODS-3 4.6 mm × 250 mm (GL Sciences, Tokyo, Japan), with a pump (JASCO, PU-2080) and a detector (JASCO, MD-2015). A gradient condition was used with eluent A ($H_2O$ with 0.1% TFA and 1% MeCN) and eluent B (MeCN with 1% $H_2O$) (A/B = 85/15 to 5/95 over 35 min). Absorbance at 254 nm was monitored.

**Determination of photostationary state (PSS) compositions (open/closed).**

A 15 mM solution of DAE620 in DMSO was irradiated with a 365 nm light (handy UV lamp; AS ONE, SLUV-4)) until the absorbance at 623 nm reached a plateau. Then, the PSS composition was determined by analytical HPLC, using the same setup mentioned above. A gradient condition was used with eluent A ($H_2O$ with 0.1% TFA and 1% MeCN) and eluent B (MeCN with 1% $H_2O$) (A/B = 85/15 to 5/95 over 35 min). Absorbance at 360 nm was monitored, which is the isosbestic point of the absorption spectra of the open and closed forms.

### Synthesis and characterization.

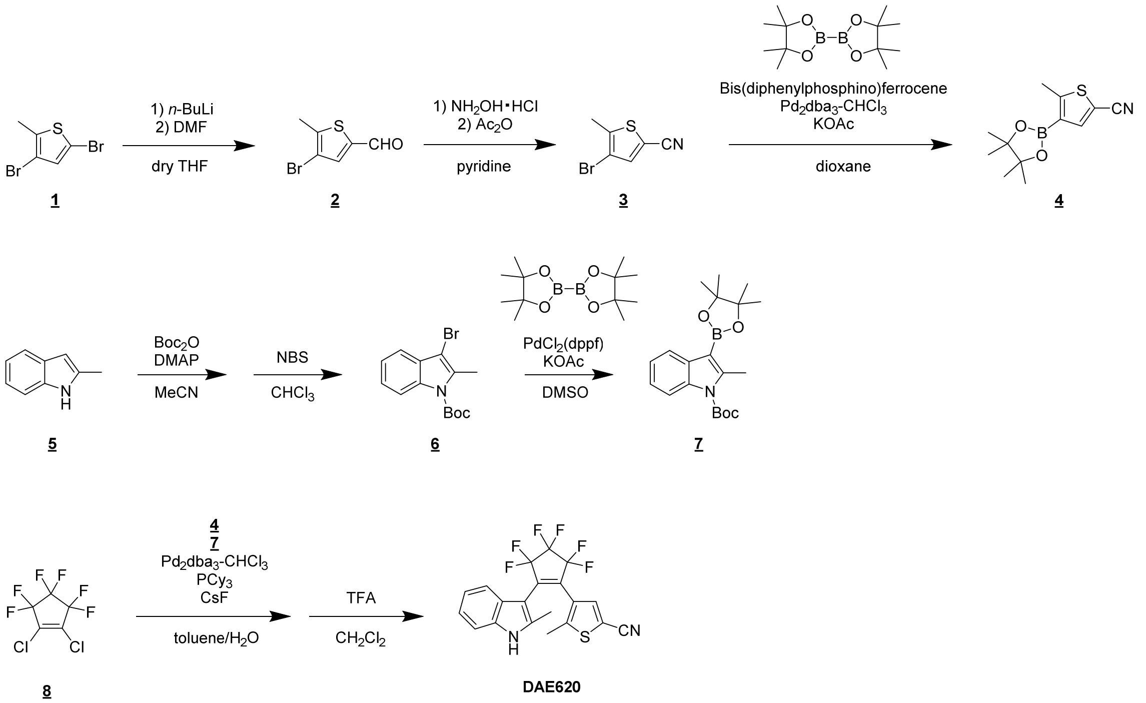


**Scheme S1.** Synthesis of DAE620.

Compound 2.

Compound 2 was synthesized according to the literature[4]. A solution of compound 1 (2.0 g, 7.8 mmol) in 3.0 mL dry THF was cooled to -78 °C under an argon atmosphere. A 1.6 M *n*-hexane solution of *n*-BuLi (5.7 mL, 9.1 mmol) was added dropwise, and the mixture was stirred for 30 min at the same temperature. Then, dry DMF (1.0 mL, 12 mmol) was slowly added. The reaction mixture was warmed to 0 °C and stirred for 1 h, then the reaction was quenched with 2 M HCl aq., and the mixture was extracted with EtOAc. The organic layer was washed with brine, dried over $Na_2SO_4$ and evaporated to dryness to give pure compound 2 (1.4 g, 84%) without further purification. $^1$H NMR (400 MHz, $CDCl_3$): δ 9.77 (s, 1H), 7.58 (s, 1H), 2.48 (s, 3H).

Compound 3.

Compound 3 was synthesized according to the literature[5]. To a solution of compound 2 (1.09 g, 5.32 mmol) in 8.0 mL pyridine was added $NH_2OH{\cdot}HCl$ (578 mg, 8.31 mmol), and the mixture was stirred for 3 h at room temperature. Then, $Ac_2O$ (1.0 mL, 10 mmol) was added, and the mixture was refluxed for 18 h at 130 °C. After cooling to room temperature, the reaction mixture was diluted with water, and extracted with $CH_2Cl_2$. The organic layer was washed with brine, dried over $Na_2SO_4$ and evaporated to dryness. The residue was purified by column chromatography (silica gel, *n*-hexane/$CH_2Cl_2$ = 100/0 to 40/60) to give compound 3 (858 mg, 80%). $^1$H NMR (400 MHz, $CDCl_3$): δ 7.44 (s, 1H), 2.46 (s, 3H); $^{13}$C NMR (101 MHz, $CDCl_3$): δ 142.84, 139.75, 113.39, 110.44, 107.49, 15.34.

Compound 4.

Compound 4 was synthesized according to the literature[5]. Compound 3 (641 mg, 3.17 mmol), bis(pinacolato)diboron (950 mg, 3.74 mmol), 1,1'-bis(diphenylphosphino)ferrocene (17.6 mg, 0.0317 mmol), $Pd_2dba_3 \cdot CHCl_3$ (16.4 mg, 0.0159 mmol) and KOAc (770 mg, 7.85 mmol) were dissolved in 6.0 mL dry 1,4-dioxane under an argon atmosphere. The mixture was refluxed for 18 h at 130 °C. After cooling to room temperature, the reaction mixture was filtered through Celite. After evaporation, the residue was purified by column chromatography (silica gel, *n*-hexane/ EtOAc = 100/0 to 77/23) to give compound 5 (442 mg, 56%). $^1H$ NMR (400 MHz, $CDCl_3$): δ 7.74 (s, 1H), 2.71 (s, 3H), 1.32 (s, 12H); HRMS ($ESI^+$): Calcd for $[M+Na]^+$, 272.08892, Found, 272.08970 (-0.8 mDa).

Compound 6.

Compound 6 was synthesized according to the literature[4]. Compound 5 (5.0 g, 38 mmol), $Boc_2O$ (9.0 g, 41 mmol) and DMAP (502 mg, 4.11 mmol) were dissolved in 30 mL MeCN, and the mixture was stirred for 18 h at room temperature. The reaction mixture was evaporated and the residue was extracted with $CH_2Cl_2$. The organic layer was washed with brine, dried over $Na_2SO_4$ and evaporated to dryness. The crude intermediate was dissolved in 20 mL $CH_2Cl_2$. To the resulting solution was added *N*-bromosuccinimide (7.5 g, 42 mmol). The mixture was stirred for 40 min at room temperature, then diluted with water, and extracted with $CH_2Cl_2$. The organic layer was washed with brine, dried over $Na_2SO_4$ and evaporated to dryness. The residue was purified by column chromatography (silica gel, *n*-hexane/$CH_2Cl_2$ = 100/0 to 54/46) to give compound 6 (9.3 g, 79%). $^1H$ NMR (400 MHz, $CDCl_3$): δ 8.12–8.08 (m, 1H), 7.48–7.43 (m, 1H), 7.31–7.26 (m, 2H), 2.64 (s, 3H), 1.68 (s, 9H).

Compound 7.

Compound 6 (2.1 g, 6.8 mmol), bis(pinacolato)diboron (2.1 g, 8.2 mmol), $PdCl_2(dppf)$ (1.0 g, 20%mol) and KOAc (2.0 g, 21 mmol) were dissolved in 20 mL dry DMSO, and the mixture was stirred for 20 h at 85 °C. After cooling to room temperature, the reaction mixture was filtered through Celite. After evaporation, the residue was purified by column chromatography (silica gel, *n*-hexane/$CH_2Cl_2$ = 70/30 to 10/90) to give compound 7 (564 mg, 23%). $^1H$ NMR (400 MHz, $CDCl_3$): δ 8.07–8.01 (m, 1H), 8.00–7.95 (m, 1H), 7.23–7.19 (m, 2H), 2.84 (s, 3H), 1.68 (s, 9H), 1.36 (s, 12H); HRMS ($ESI^+$): Calcd for $[M+Na]^+$, 380.20072, Found, 380.20176 (-1.0 mDa).

DAE620.

Compound 8 (696 mg, 1.95 mmol), $Pd_2dba_3 \cdot CHCl_3$ (102 mg, 0.0985 mmol), $PCy_3$ (106 mg, 0.378 mmol) and CsF (2.4 g, 16 mmol) were dissolved in 3.0 mL toluene and 0.9 mL water under an argon atmosphere. Compound 4 (442 mg, 1.77 mmol) in 3.0 mL toluene and compound 7 (696 mg, 1.95 mmol) in 3.0 mL toluene were added, and the mixture was refluxed at 120 °C for 18 h. After cooling to room temperature, the reaction mixture was filtered through Celite. The filtrate was evaporated and the residue was roughly purified by column chromatography (silica gel, *n*-hexane/$CH_2Cl_2$ = 100/0 to 0/100). The crude intermediate was dissolved in 1.0 mL $CH_2Cl_2$, and then 1.0 mL TFA was added. The mixture was stirred for 1 h at room temperature, then quenched with sat. $NaHCO_3$ aq. and extracted with $CH_2Cl_2$. The organic layer was washed with brine, dried over $Na_2SO_4$ and evaporated to dryness. The residue was purified by column chromatography (silica gel, *n*-hexane/$CH_2Cl_2$ = 46/54 to 0/100) to give DAE620 (32 mg, 4%). $^1H$ NMR (400 MHz, $CDCl_3$): δ 8.22 (brs, 1H), 7.68 (s, 1H), 7.41 (d, $J$ = 8.0 Hz, 1H), 7.31 (d, $J$ = 8.0 Hz, 1H), 7.20 (ddd, $J$ = 8.1, 7.6, 1.2 Hz, 1H), 7.12 (ddd,

$J$ = 8.2, 7.6, 1.1 Hz, 1H), 2.08 (s, 3H), 1.86 (s, 3H); $^{13}$C NMR (101 MHz, $CDCl_3$): δ 148.59, 137.78, 136.42, 135.31, 126.81, 125.91, 122.88, 121.53, 119.25, 113.43, 110.82, 107.63, 101.25, 14.76, 13.10; HRMS (ESI$^-$): Calcd for $[M\text{-}H]^-$, 425.05526, Found, 425.05740 (-2.1 mDa). HPLC analysis: peak at 22.2 minutes corresponds to DAE620.

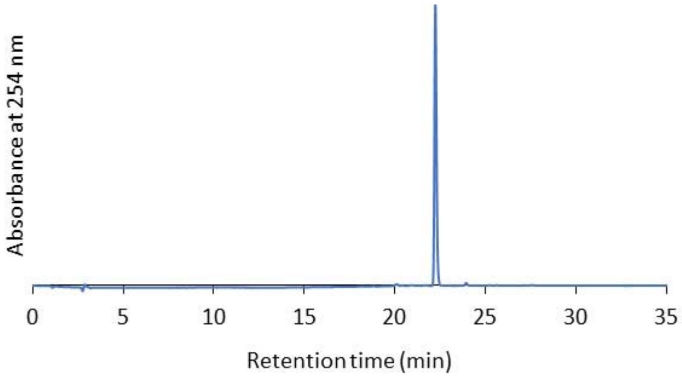

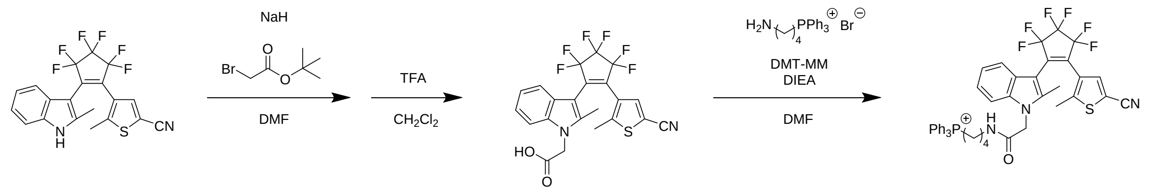


**DAE620** **9** **DAE620-Mito**

**Scheme S2.** Synthesis of DAE620-Mito.

Compound 9.

A solution of DAE620 (17.7 mg, 0.042 mmol) in 0.8 mL dry DMF was cooled to 0 °C under an argon atmosphere. NaH (12.0 mg, 0.50 mmol) was added slowly, and the mixture was stirred for 30 min at the same temperature. Then, *tert*-butyl bromoacetate (64.7 mg, 0.33 mmol) was added. The reaction mixture was warmed to room temperature and stirred for 18 h, then the reaction was quenched with water and the mixture was extracted with EtOAc. The organic layer was washed with brine, dried over $Na_2SO_4$ and evaporated to dryness. The residue was roughly purified by column chromatography (silica gel, *n*-hexane/EtOAc = 83/17 to 62/38). The crude intermediate was dissolved in 0.5 mL $CH_2Cl_2$, and then 1.0 mL TFA was added. The mixture was stirred for 1 h at room temperature, then quenched with sat. $NaHCO_3$ aq. and extracted with $CH_2Cl_2$. The organic layer was washed with brine, dried over $Na_2SO_4$ and evaporated to dryness to give pure compound 9 (12.5 mg, 62%) without further purification. $^1$H NMR (400 MHz, $CD_3OD$): δ 7.79 (s, 1H), 7.34 (d, *J* = 8.0 Hz, 1H), 7.26 (d *J* = 8.2 Hz, 1H), 7.11 (ddd, *J* = 8.3, 7.1, 1.2 Hz, 1H), 7.01 (ddd, *J* = 8.1, 7.1, 1.0 Hz, 1H), 4.86 (s, 2H), 1.88 (s, 3H), 1.69 (s, 3H); $^{13}$C NMR (101 MHz, $CD_3OD$): δ 171.39, 151.12, 140.41, 139.17, 138.69, 127.79, 126.48, 123.66, 122.46, 119.95, 114.21, 110.58, 108.72, 102.02, 45.56, 14.95, 11.59; HRMS ($ESI^-$): Calcd for $[M-H]^-$, 483.06074, Found, 483.06144 (-0.7 mDa).

DAE620-Mito.

A solution of compound 9 (2.92 mg, 0.0060 mmol) in 0.2 mL DMF was cooled to 0 °C. Then, (4-ammoniobutyl)triphenylphosphonium bromide (13.7 mg, 0.028 mmol, synthesized according to the reported procedure[6]), DMT-MM (10.0 mg, 0.036 mmol) and DIEA (18.5 mg, 0.14 mmol) were added, and the mixture was stirred for 30 min at the same temperature. After warming to room temperature, the mixture was further stirred for 18 h, and then evaporated. The residue was purified by preparative HPLC using eluent A ($H_2O$ with 0.1% TFA) and eluent B (MeCN) (A/B = 90/10 to 0/100 over 40 min) to give DAE620-Mito (3.17 mg, 66%). $^1$H NMR ($CD_3OD$, 400 MHz, partially overlapped with solvent peaks): δ 7.78–7.72 (m, 4H), 7.67–7.54 (m, 12H), 7.37–7.34 (m, 1H), 7.14–7.10 (m, 1H), 7.04–6.98 (m, 2H), 4.69 (s, 2H), 3.38–3.29 (m, 2H), 3.24–3.15 (m, 2H), 1.87 (s, 3H), 1.71 (s, 3H), 1.66–1.61 (m, 2H), 1.59–1.53 (m, 2H); HRMS ($ESI^+$): Calcd for $[M]^+$, 800.22937, Found, 800.23020 (-0.8 mDa). HPLC analysis: peak at 18.3 minutes corresponds to DAE620-Mito.

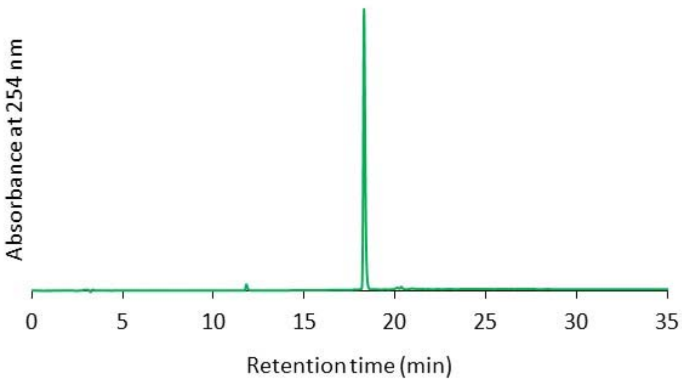
Absorbance at 254 nm
0
5
10
15
20
25
30
35
Retention time (min)

### Supporting References


1. Irie, M. Diarylethenes for memories and switches. *Chem. Rev.* **100**, 1685–1716 (2000).
2. Frisch, M. J., Trucks, G. W., Schlegel, H. B., Scuseria, G. E., Robb, M. A., Cheeseman, J. R., Scalmani, G., Barone, V., Petersson, G. A., Nakatsuji, H., Li, X., Caricato, M., Marenich, A., Bloino, J., Janesko, B. G., Gomperts, R., Mennucci, B., Hratchian, H. P., Ortiz, J. V., Izmaylov, A. F., Sonnenberg, J. L., Williams-Young, D., Ding, F., Lipparini, F., Egidi, F., Goings, J., Peng, B., Petrone, A., Henderson, T., Ranasinghe, D., Zakrzewski, V. G., Gao, J., Rega, N., Zheng, G., Liang, W., Hada, M., Ehara, M., Toyota, K., Fukuda, R., Hasegawa, J., Ishida, M., Nakajima, T., Honda, Y., Kitao, O., Nakai, H., Vreven, T., Throssell, K., Montgomery, J. A., Jr., Peralta, J. E., Ogliaro, F., Bearpark, M., Heyd, J. J., Brothers, E., Kudin, K. N., Staroverov, V. N., Keith, T., Kobayashi, R., Normand, J., Raghavachari, K., Rendell, A., Burant, J. C., Iyengar, S. S., Tomasi, J., Cossi, M., Millam, J. M., Klene, M., Adamo, C., Cammi, R., Ochterski, J. W., Martin, R. L., Morokuma, K., Farkas, O., Foresman, J. B. & Fox, D. J. *Gaussian 09*, Revision E.01, Gaussian, Inc., Wallingford CT (2016).
3. Scott, A. & Radom, L. Harmonic vibrational frequencies: An evaluation of Hartree−Fock, Møller−Plesset, quadratic configuration interaction, density functional theory, and semiempirical scale factors. *J. Phys. Chem.* **100**, 16502–16513 (1996).
4. Fredrich, S., Bonasera, A., Valderrey, V. & Hecht, S. Sensitive assays by nucleophile-induced rearrangement of photoactivated diarylethenes. *J. Am. Chem. Soc.* **140**, 6432–6440 (2018).
5. Hiroto, S., Suzuki, K., Kamiya, H. & Shinokubo, H. Synthetic protocol for diarylethenes through Suzuki-Miyaura coupling. *Chem. Commun.* **47**, 7149–7151 (2011).
6. Duca, M., Dozza, B., Lucarelli, E., Santi, S., Di Giorgio, A. & Barbarella, G. Fluorescent labeling of human mesenchymal stem cells by thiophene fluorophores conjugated to a lipophilic carrier. *Chem. Commun.* **46**, 7948–7950 (2010).
